# Examining the relationships between phenotypic plasticity and local environments with genomic structural equation models

**DOI:** 10.1101/2019.12.11.873257

**Authors:** Malachy T. Campbell, Haipeng Yu, Mehdi Momen, Gota Morota

**Author notes:** **Corresponding Author:** Malachy T. Campbell, 417 Bradfield Hall, Cornell University, Ithaca, New York 14853.

## Abstract

Environmental association analyses (EAA) seek to identify genetic variants associated with local adaptation by regressing local environmental conditions at collection sites on genome-wide polymorphisms. The rationale is that environmental conditions impose selective pressure on trait(s), and these traits are regulated in part by variation at a genomic level. Here, we present an alternative multivariate genomic approach that can be utilized when both phenotypic and environmental data are available for the population. This framework utilizes Bayesian networks (BN) to elucidate interdependancies between local environmental conditions and empirical phenotypes, and jointly estimates the direct and indirect genetic covariances between empirical phenotypes and environmental conditions using a mixed-effects structural equation model (SEM). Direct genomic covariance between empirical phenotypes and environmental conditions may provide insight into whether QTL that affect adaptation to an environmental gradient also affects the observed phenotype. To demonstrate the utility of this approach, we leveraged two existing datasets consisting of 55 climate variables for 1,130 *Arabidopsis* accessions and empirical phenotypes for fitness and phenology collected on 515 accessions in two common garden locations in Europe. BN showed that plasticity for fitness and phenology was highly dependant on local environmental conditions. Moreover, genomic SEM revealed relatively high positive genomic correlation between plasticity in fitness and environmental variables that describe the favorability of the local environment for plant growth, indicating the presence of common QTL or independent QTL that are tightly linked. We believe the frameworks presented in this manuscript can provide new insights into the genetic basis of local adaptation.

## Introduction

Identifying traits that confer adaptation to a given environment and elucidating the genetic determinants driving variation for these traits is an important goal for physiologists, evolutionary biologists, and quantitative geneticists. In many cases, particularly those working with agronomic species, these studies involve large-scale phenotypic evaluations in multiple environments, which are later integrated with genomic data using quantitative genetic frameworks. However, when the population is composed of individuals sampled across an environmental gradient, information regarding local environmental conditions at collection sites can be leveraged together with genomic data to identify genetic variants associated with variation for a given environmental factor (Fournier-Level et al., 2011; Blanquart et al.,2013; Yoder et al., 2014; Tiffin and Ross-Ibarra, 2014; Hoban et al., 2016). In recent years, a number of studies have employed similar approaches, termed environmental association analysis (EAA), to study the genetic basis of local adaptation (Fournier-Level et al., 2011; Yoder et al., 2014; Lasky et al., 2015).

EAA seeks to identify genomic variants that are associated with variation in environmental conditions at collection sites (Jones et al., 2013; Dell’Acqua et al., 2014; Yoder et al., 2014; Lasky et al.,2015; Anderson et al., 2016). The rationale for these approaches is that local environmental conditions impose selective pressure on some trait(s), and these traits are regulated in part by variation at a genomic level. Since adaptive traits should be correlated with local environmental conditions, regression of environmental variables on genome-wide single nucleotide polymorphisms (SNPs) may yield markers that are associated with environmental variables and, by proxy, adaptive traits. Several studies have leveraged these, and similar approaches, to elucidate the genetic basis of local adaptation (Fournier-Level et al., 2011; Yoder et al., 2014; Lasky et al., 2015). The only requirements for EAA is genomic data for a georeferenced population and environmental variables recorded at, or close to, collection sites. Downstream analyses or independent studies are performed to determine if these variants have an effect on the phenotype, or whether they can be used to predict phenotypic variation. For instance, Yoder et al.(2014) utilized a population of 202 wild *Medicago truncatula* accessions to identify genomic associations with annual mean temperature, precipitation in the wettest month, and isothermality. They showed that accessions with a greater number of alleles associated with high precipitation in the wettest month also exhibited higher growth rate in a wet controlled environment. Similarly, Lasky et al. (2015) first identified environment-genotype associations in a panel of *Sorghum* landraces, and used these associations to predict agronomic characteristics in environments with contrasting moisture or edaphic conditions. Thus, these studies provide evidence that EAA can recover genetic determinate that are associated with environmental adaptation, and may influence phenotypic variation for adaptive or agronomically relevant traits.

However, when both phenotypic and environmental data are available for the population, alternative multivariate approaches can be utilized to jointly estimate genomic parameters and elucidate the genetic interrelationships between local environmental conditions and observable phenotypes. With these approaches we can address whether there is a dependancy between the empirical phenotype and the local environmental condition, effectively addressing the question “Is local adaptation to an environmental variable dependant on this trait?” and “What genes have an impact on both local adaptation *and* the empirical phenotype?” Structural equation models (SEM) are powerful frameworks that can be used to model the interdependancies between multiple variables (Wright, 1921; Haavelmo, 1943). When integrated into a quantitative genetics framework, these approaches allow quantitative genetic loci (QTL) or total genomic values to be decomposed into direct and indirect effects based on a predefined graphical model that describes directed relationships between variables (Gianola and Sorensen, 2004; Valente et al.,2013). SEM can be viewed as an extension of a conventional multi-trait (MT) quantitative genetic framework (Valente et al., 2013). Whereas with MT approaches, covariances among observable phenotypes are estimated and used to describe the symmetric linear relationships between variables, SEM extends the multivariate framework to allow recursive (effects from one phenotype affects the outcome of another) and simultaneous (reciprocal) structures among its variables by utilizing phenotypes as predictors for other phenotypes (Goldberger, 1972; Bielby and Hauser, 1977).

In quantitative genetics, SEM has been largely applied to topics in animal breeding and genetics. For instance in one of the first applications of SEM in quantitative genetics in the context of a linear mixed model, de los Campos et al. (2006b) used SEM to elucidate the interrelationship between milk yield and mastitis (inflammation of the udder quantified using somatic cell scores) in dairy cattle. The authors showed that models where milk yield was dependant on mastitis were better supported by the data, indicating that disease was the primary driver of reduced milk production rather than the converse. Since this work, quantitative genetic SEM frameworks have been used to elucidate the genetic interdependencies among meat quality traits, calving traits, fertility metrics, as well as milk yield and mastitis in other species or breeds (de los Campos et al., 2006a,b; Varona et al., 2007; Wu et al., 2007; König et al., 2008; Heringstad et al., 2009; de Maturana et al., 2009, 2010; Jamrozik et al., 2010; Peñagaricano et al., 2015a,b). More recently, the SEM quantitative genetic frameworks have been extended to perform genome-wide associations in chicken and rice (Momen et al., 2018, 2019). Given that many EAA studies assume a causal relationship between an unobserved phenotype and the local environment, SEM provides a framework where in these relationships can be explicitly encoded in the model – when empirical phenotypes are available for the same population. Moreover, these frameworks provide a means to examine the covariance in genetic effects that act directly on the empirical phenotype and the environmental variable (Valente et al., 2013).

Direct applications of quantitative genetic SEM frameworks to EAA is not trivial. For one, SEM requires a putative causal networks that describes the dependancies among and between environmental variables and empirical phenotypes (Gianola and Sorensen, 2004). In most cases, these networks are not only unknown, but learning the structure may even be an objective of the study itself. Secondly, the environmental data often consist of dozens or hundreds of variables that are highly correlated (Ferrero-Serrano and Assmann, 2019; Lasky et al., 2015). Thus, prior to applying SEM to EAA we must reduce the dimensionality of the environmental data and determine an appropriate network structure. One popular approach for dimensional reduction is factor analysis (FA) (de Los Campos and Gianola,2007). The underlying rationale for FA is that relationships among variables are due to some underlying unobserved process. The goal of FA is to define a reduced set of unobserved, latent variables that maximize the correlation among groups of related observed variables. In quantitative genetics, FA is routinely applied to multi-environmental trials and high-dimensional multi-trait applications (Kelly et al., 2007; Meyer, 2009; de Los Campos and Gianola, 2007; Runcie and Mukherjee, 2013; Yu et al.,2019). Thus, when applied to high dimensional environmental data, FA may yield a reduced set of underlying variables that capture major patterns of local environments. When the underlying causal structure is unknown, Bayesian network (BN) approaches can be utilized to elucidate the probabilistic dependencies among variables (Scutari, 2009; Scutari and Denis, 2014). These dependencies are expressed using a directed acyclic graph where each variable is depicted as a node and directed edges join dependant nodes. Although BN approaches learn dependencies from the data itself, these approaches can yield insightful information regarding the causal relationships among variables. Such approaches have been leveraged to understand the genetic interdependencies among complex traits and have been utilized to elucidate potential causal structures that can be used in SEM quantitative genetic frameworks (Valente et al., 2013; Yu et al., 2019; Momen et al., 2018, 2019). Thus, both FA and BN can be leveraged to reduce the dimensionality of local environmental variables and elucidate the relations between traits or latent factors.

The objective of this study is to demonstrate the utility of SEM quantitative genetic frameworks for studying the genetic interrelationships between local environmental conditions and empirical phenotypes associated with fitness and phenology. To this end, we utilized two publicly available data sets that describe environmental variables at collection sites for 1,130 diverse *Arabidopis thaliana* accessions, and empirical phenotypes in two precipitation regimes at two common garden locations in Europe (Ferrero-Serrano and Assmann, 2019; Exposito-Alonso et al., 2019). Several studies have shown that adaptation is polygenic (Pritchard and Di Rienzo, 2010; Pritchard et al., 2010; Flood and Hancock,2017). With this in mind, we sought to forego single marker inferences and instead predict total genomic values for each individual, which are the cumulative additive genetic value for a given phenotypic variable. We seek to decompose total genetic effects for these variables into direct and indirect effects, effectively allowing us to address the following questions: “Are genetic effects for empirical phenotypes dependant on the genetic drivers for adaptation to local environmental conditions (and vice versa)?” and “How much of the total genomic value for an empirical phenotype is due to genetic effects from upstream phenotypic variables?”.

## Materials and Methods

### Environmental variables

This study utilized a publicly available data set of local environmental conditions for 1,130 *Arabidopsis* accessions. The original data, compiled by Ferrero-Serrano and Assmann (2019), consisted of 205 environmental variables for 829 unique collection sites. Categorical variables were removed from the data set, as well as variables that had missing values in ≥ 20% of the accessions. After this filtering, 139 climate variables remained. Prior to FA, we removed variables that showed high collinearity, as variables with very high correlation can interfere with factor analysis. In total, these quality control steps provided data for 55 environmental variables for 1,130 accessions.

### Empirical Phenotypes

Since the objective of the study was to examine the genomic interrelationships between local climate conditions and phenotypic plasticity in contrasting environments, we sought a data set that provided phenotypes recorded in the same germplasm in contrasting and ecologically-relevant conditions. To this end, we used data from a recent study by Exposito-Alonso et al. (2019) in which 515 of the 1,130 accessions were phenotyped for fitness, germination time and flowering time in two locations within the natural range of *Arabidopsis thaliana* and two simulated precipitation regimes. The experimental design and collection of phenotypic data is explained in great detail by Exposito-Alonso et al. (2019). Briefly, the 515 accessions were grown in open-ended rain-out shelters in Tuebingen, Germany and Madrid, Spain. The open-ended design allows for the temperature and humidity conditions within the structure to be similar to the natural environment. Within each location the plants were grown in a split-plot design. Two simulated precipitation regimes, which were designed to mimic natural rainfall at Tuebingen and Madrid, were randomly assigned to each subplot. The interquartile range for soil water content (SWC) in the low-precipitation treatment was 11.38-22.51% with a median of 16.1% in Madrid and 10.76-20.09% with a median of 14.7% in Tuebingen. The interquartile range for the high precipitation regime was 20.73-29.02% with a median of 24.6% in Madrid, and 22.62-33.00% with a median of 27.8% in Tuebingen. Median midday photosynthetically active radiation (PAR) values inside the shelters were 45.7 mol·m^−2^·day^−1^ in Madrid and 30.9 mol·m^−2^·day^−1^ in Tuebingen. Temperatures outside the structures ranged from 5.34-12.39°C with a median of 8.5°C in Madrid and 2.44-9.54°C with a median of 5.6°C in Tuebingen. These ranges are very consistent with temperatures recorded in the structures (Exposito-Alonso et al., 2019).

We estimated the macroenvironmental sensitivity for each accession and each empirical phenotype that was recorded by Exposito-Alonso et al. (2019) using the Finlay-Wilkinson (FW) approach (Finlay and Wilkinson, 1963). FW essentially expresses the plasticity of an accession grown across multiple environments as a function of the overall population performance in each environment. The FW model is given by

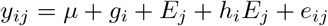

where *y_ij_* is the phenotype for accession *i* in environment *j, μ* is the overall mean, *g_i_* is the main accession effect, *E_j_* is the main environment effect, *h_i_* is the slope for accession *i* on the overall environment means, and *e_ij_* is the residual for accession *i* in environment *j*. Here, *y_ij_* are best linear unbiased estimates for the accession effect in each environment from a model that accounts for systematic experimental effects (Exposito-Alonso et al., 2019). The FW model was fit using the `FW` package in R (Lian, 2014). The slope from this model was used as a metric for phenotypic plasticity in all downstream analysis.

### Genotyping data

Imputed SNP markers were obtained for all 1,135 accessions from 1001genomes (https://1001genomes.org/data/GMI-MPI/releases/v3.1/SNP_matrix_imputed_hdf5/)(Weigel and Mott, 2009; Alonso-Blanco et al., 2016). We extracted marker information for the 1,130 accessions with climate data, and removed SNPs with low minor allele frequencies (MAF < 0.05). Moreover, SNPs in high linkage disequilibrium (LD) (*r* > 0.85) were removed using the PLINK indep function with a 50 SNP window, a step size of 5 SNPs, and a variance inflation factor (VIF) of 3.6. The VIF is computed as 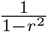. Thus, a VIF of 3.6 corresponds to a *r* ≈ 0.85. After these filtering steps, 426,567 SNPs remained.

### Factor analysis of environmental variables

To reduce the dimensionality of the 55 environmental variables, and define a reduced subset that captures potential undefined/unobserved variables that give rise to the original covariance, we utilized a combination of FA techniques, specifically exploratory and confirmatory factor analysis (EFA and CFA, respectively). Factor analysis seeks to identify a smaller set of latent variables that capture the underlying interrelationships between the original, manifest variables. The relationships between latent and manifest variables is given by

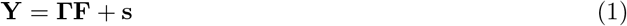

where **Y** is an *t* × *n* matrix of phenotypes with *n* = 515 indicating the number of accessions and *t* = 55 indicating the number of traits; **F** is an *l* × *n* matrix of factor scores that describe the values for each latent factor (*l*) for each accession; **Γ** is an *t* × *l* matrix that shows how each trait (*t*) loads onto each latent factor; and s is a *t* × *n* matrix that represents the specific effects for each trait and accession. Thus, FA expresses a set of manifest variables as a function of common, latent factors.

While both EFA and CFA are based on a similar framework, EFA allows manifest variables to load onto multiple latent factors and CFA does not. Thus, EFA is most often used to determine the appropriate number of latent factors and examine how manifest variables load on to them, and CFA is used to test hypothesis regarding the relationships between manifest and latent factors and to estimate factor loading scores. We determined the appropriate number of factors using parallel analysis. Parallel analysis is a simulation-based method that was originally proposed by Horn (1965) to determine the optimal number of latent factors. Briefly, parallel analysis randomly simulates data sets with similar properties to the observed data and uses these data to extract eigenvalues. Scree plots are used to plot and compare eigenvalues from the simulated data and eigenvalues from the observed data. The optimal number of factors is determined as the maximum number of factors that have observed eigenvalues that are larger than eigenvalues from simulated data. Parallel analysis was performed using the `fa.parallel` function in the `psych` package R (Revelle, 2018). We used the minimum residual method with 1,000 iterations. Once the optimal number of factors was determined (11 latent factors), EFA was performed using the factor analysis function, `fa()`, with varimax rotation and the minimum residual method with 1,000 iterations.

CFA was used to estimate factor scores for each accession and latent environmental variable. Since CFA only allows manifest variables to load onto a single latent variable, we used EFA results to determine which latent factor had the largest absolute loading for each manifest variable. Although EFA identified 11 latent factors, one latent factor was omitted from CFA because all manifest variables that loaded onto this latent factor had higher loadings for other latent factors. CFA was fit using the `sem` package in R according to the loadings provided in Figure 1 (Fox et al., 2017). Factor scores were computed with the ‘regression’ method using the `fscores()` function in the `sem` package (Fox et al., 2017).

**Figure 1.**
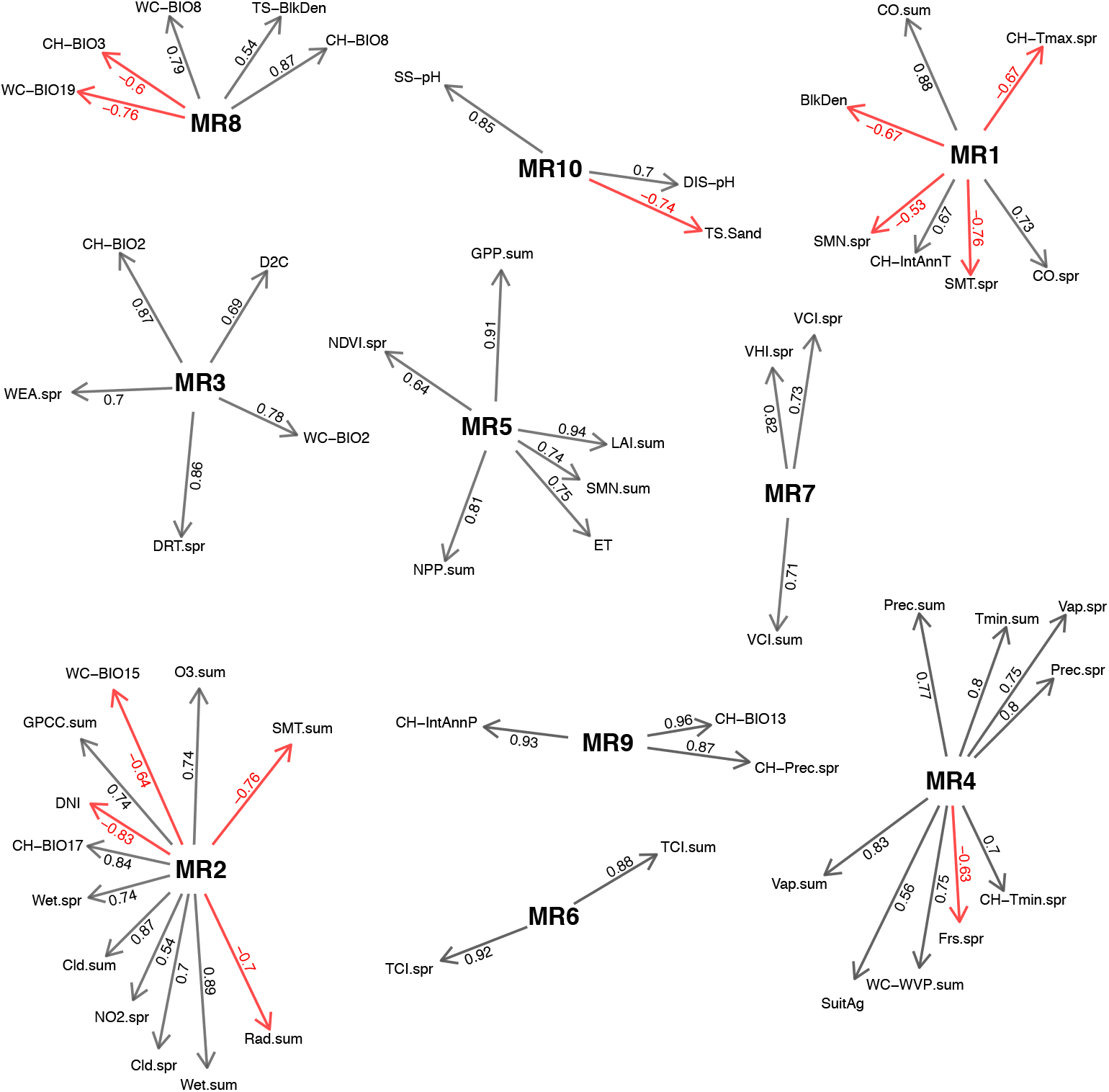
Factor loadings for manifest local environmental variables. Variables in bold type face are latent factors identified using factor analysis, while nodes emanating from these are manifest environmental variables. Edges colored in grey indicate the manifest variable has a positive loading on the latent factor, while those in red indicate negative loadings.

### Structure learning using Bayesian network

We next sought to elucidate the genomic interrelationships between plasticity and latent factor scores from CFA for local environmental conditions following an approach described by Yu et al. (2019). To this end, we first predicted genomic values for each accession and trait using a Bayesian multi-trait model (MTM). The MTM is given by

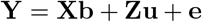

where **Y** is an *n* × *t*′ matrix of phenotypes composed of factor scores for latent environmental factors and plasticity for empirical phenotypes (*t*′ = 13), where *n* = 515 is the number of individuals and *t*′ is the number of phenotypes (ten latent local environmental variables and three empirical phenotypes, *t*′ = 13); **X** and **Z** are incidence matrices that relate phenotypes to vectors of systematic effects (**b**) and additive genetic effects **u**, respectively; and **e** is the error term. Moreover, we assume **u** ~ N(0, **Σ_u_** ⊗ **G**) and **e** ~ N(0, **Σ_e_** ⊗ **I**_*n*×*n*_), where **G** is a genomic relationship matrix constructed following VanRaden (2008), **Σ_u_** is a *t*′ × *t*′ covariance matrix for additive genetic effects. The `MTM` was fit using the MTM package in R with 10,000 Markov chain Monte-carlo (MCMC) samples of which the first 2,000 are discarded and every fifth sample was retained (de los Campos and Grüneberg, 2016).

Bayesian network (BN) learning approaches assume that the samples are independent. However, when predicting additive genomic values using MTM, dependencies are between breeding values for accessions are introduced from **G**. Therefore prior to BN learning, we followed an approach described by Töpner et al. (2017) to remove dependencies. Briefly, **G** was decomposed into Cholesky factors by **G** = **LL′**, where **L** is a lower triangle matrix with dimensions *n* × *n*. We define a *nt*′ × *nt*′ matrix, **M** via **M** = **I**_*t*′ × *t*′_ ⊗ **L**. By multiplying the *nt*′ vector of genomic values (**u**) by the inverse of **M**, we are provided with a vector of transformed genomic values (**u**\* = **M**^−1^**u**) that follow a distribution given by N(0, **Σ_*g*_** ⊗ **I**_*n* × *n*_). Thus, the transformed genomic values are independent between accessions.

BN are a class of graphical models that represent the probabilistic dependencies between a set of random variables as a directed acyclic graph 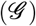 (Scutari and Denis, 2014). 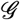 is composed of nodes (*V*) that represent random variables and edges (*E*) that depict probabilistic dependencies between nodes. BN follow the Markov property, which states that given its parents, a node is conditionally independent of all nodes that are non-descendants (Scutari and Denis, 2014). The joint probability distribution for *k* random variables (*X_v_* = (*X*_1_,…, *X_k_*)) is given by

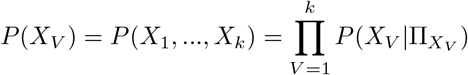

where parent nodes to *X_v_* is indicated by Π_*X_V_*_ (Scutari and Denis, 2014).

The vector of transformed genomic values (**u**\*) was used as input for BN learning using the `bnlearn` package (Scutari, 2009). Structure learning was performed using four algorithms: hill-climbing (HC), tabu-search, max-min hill-climbing (MMHC), and general 2-phase restricted maximization (RSmax2). HC and tabu are score-based, greedy algorithms which seek to maximize the goodness-of-fit (i.e., network score). These algorithms begin with an empty network structure and add, remove, or reverse edge each edge until a maximum score is reached. The latter two algorithms, MMHC and RSmax2, are hybrid learning algorithms, which essentially restrict the score-based approach described above on a subset of nodes within the network (Tsamardinos et al., 2006). For each algorithm, we used a combination of bootstrapping and model averaging to identity robust networks and quantify uncertainty in linkages and the direction of each edge. Five hundred bootstrapping replicates were used and edges that were present in less than 85% of the networks were removed, and the models were averaged. We compared networks from each algorithm using the Bayesian information criteria (BIC) and selected the ‘best’ network according to the network that produced the highest BIC since `BNlearn` rescales BIC values by −2.

### Genomic structural equation model

Work by Gianola and Sorensen (2004) provided a basis to introduce SEM into classical quantitative genetics frameworks. SEM utilize a system of linear equations to model the interrelationships between multiple dependant variables. Once introduced into the quantitative genetics frameworks pioneered by Henderson (1984), these approaches provide a means to partition multiple phenotypes into direct and indirect genetic components according to a predefined network structure (Gianola and Sorensen, 2004;Valente et al., 2013; Bello et al., 2018). In matrix form, the structural equation model is given by

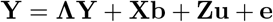

where all matrices are defined according to the MTM described above. However, note that the response variable **Y** appears on both the right and left-hand side of the equation, meaning that some phenotypes will serve as covariates for other phenotypes. The effect of an upstream phenotype on a downstream phenotype is determined by the direction and magnitude of elements in the coefficient matrix (**Λ**). **Λ** is typically a lower triangle matrix with zeros in the diagonal and upper triangle. We assume **u** ~ N(0, **Σ_*u*_0__**) ⊗ **G**) and **e** ~ N(0, **Σ_*e*_0__** ⊗ **I**_*n* × *n*_), where **Σ_*u*_0__** and **Σ_*e*_0__** represent the genomic and residual covariances for total effects.

Given a simple, hypothetical causal structure for three phenotypes (*y*_1_ → *y*_2_ and *y*_1_ → *y*_3_), we can decompose each phenotype into genetic and non-genetic components using the following system of equations

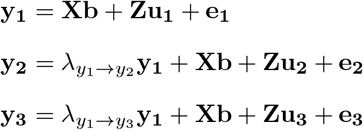

Since *y*_1_ has no variables leading to it, the total genomic effects for *y*_1_ are given by **u_1_total__** = **u_1_**. For *y*_2_ we have an indirect effect coming from *y*_1_, therefore the total genomic value is given by **u_2_total__** = λ_*y*_1_ → *y*_2__ **u**_1_ + **u_2_**. For *y*_3_, total genomic values are given by **u_3_total__** = λ_*y*_1_ → *y*_3__ **u_1_** + **u_3_**. Solving the mixed model equation provides solutions for direct genomic values and estimates the genetic and residual (co)variances for direct effects among traits (**Σ_u_0__** and **Σ_e_0__**, respectively). Covariances for total genomic and residual effects can be computed through a simple transformation on the appropriate covariances matrix for direct effects. The total genomic covariance is given by **Σ_*g*_** = (**I**_*t*×*t*_ − **Λ**)^−1^ **Σ_u_0__**(**I**_*t* × *t*_ → **Λ**)^−1^′. We fit SEM using the ten latent environmental variables and the plasticity measures for three empirical phenotypes according to the learned structure described above. The model was fit using the `MTM` package with 10,000 MCMC samples with the first 2,000 samples discarded and every fifth sample retained (de los Campos and Grüneberg, 2016).

### Data availability

Local environmental variables were obtained from the Arabidopsis ClimTools repository (https://github.com/CLIMtools)(Ferrero-Serrano and Assmann, 2019), and empirical phenotypes for common garden locations were obtained from Exposito-Alonso et al. (2019). Scripts used for analyses of these data are available Arabidopsis EFA repository (https://github.com/malachycampbell/ArabidopsisEFA) and are documented to ensure reproducibility. Supplemental figures and files are available at FigShare ().

## Results

To examine the genomic relationship between local environments across the native range of *Arabidopsis thaliana* we utilized a publicly available panel of 1,135 diverse *Arabidopsis* accessions. These materials were collected from 829 non-redundant sites across Europe, Asia, Africa and N. America, and are discussed in detail by Ferrero-Serrano and Assmann (2019). The collection site for each accession is provided as Supplemental Figure S1. We utilized an existing dataset of 205 climatic, edaphic, and remote sensing variables to characterize the local environmental conditions at each of the collection sites. These variables describe precipitation, temperature, and vegetative productivity patterns, as well as soil physical and chemical characteristics (Ferrero-Serrano and Assmann, 2019).

### Factor analysis reveals the underlying structure of local environments

An initial inspection of the environmental variables showed a high degree of correlation between variables (Supplemental Fig. S2). Given the size of the data set, as well as the high degree of correlation between variables, we sought to reduce the 55 variables to a smaller set of factors that capture the underlying theoretical structure of the environments. To this end, we performed EFA on the set of 55 variables to explore the underlying structure of local environmental conditions and define a reduced set of variables that capture unobserved processes (latent factors) that drive these relationships. Confirmatory factor analysis was used to determine the contribution of each environmental variable to the latent factor and quantify how each accession contributed to each latent factor. EFA revealed that the 55 variables could be reduced to a set of 11 latent factors (Supplemental Fig. S3). Although 11 latent factors were defined, variables loading onto factor 11 had stronger loading on other latent factors. Thus this latent factor was omitted from downstream analysis. The loadings from EFA are provided as Supplemental File S1.

In theory, these latent factors should represent unobserved processes that give rise to the observed variables, and in the context of the current study, may describe processes that shape local environments. Factor loadings from CFA are shown in Figure 1. A complete listing of latent factors, the manifest variables that load onto them, and the interpretation of latent factors is provided in Supplemental File S2. Twelve environmental variables loaded onto the second latent factor (MR2). The manifest variables describe the frequency of wet days, cloud coverage, solar radiation, precipitation seasonality and precipitation of the driest quarter. Variables associated with precipitation and cloud cover largely showed positive contributions to MR2, while those associated with solar radiation showed negative contributions. Thus, MR2 is likely a description of how bright and dry an environment is. Three latent factors were defined which captured the favorability of local environments to plant growth. For instance, two metrics for vegetation condition index (VCI) which quantifies vegetation cover in a period of time to relative extremes and vegetative health index (VHI) that represents the favorability of the environment for vegetation activity showed positive loadings onto onto MR7. Moreover, the two manifest variables that represent temperature condition index (TCI) which loaded positively onto MR6. While MR6 and MR7 are largely associated with indices that describe the potential impact of environmental conditions on plant health, MR5 captures the productively of the environment as manifest variables associated with gross primary productivity, evapotranspiration, normalized difference vegetation index, and net primary productivity were loaded onto this latent factor. Several other latent factors were identified that captured precipitation patterns at each local environment. For instance, the ninth latent factor (MR9) largely captures precipitation and precipitation variability between years. Environmental variables representing the amount of precipitation in the wettest month, precipitation in the spring, and interannual precipitation showed strong positive contributions to MR9.

### Examining plasticity in fitness and phenology in contrasting environments

The ability of plants to exhibit plasticity in phenotypic traits is important strategy for adaptation to environmental constraints. With this in mind, we sought to elucidate the genetic interrelationships between plasticity in phenological traits and fitness, and local environmental characteristics. We utilized an existing dataset consisting of phenological (time to germination and flowering) traits and fitness recorded on 515 diverse Arabidopisis accessions grown in common garden experiments in Tuebingen and Madrid (Exposito-Alonso et al., 2019). At each common garden location, accessions were grown under simulated high and low rainfall conditions, with high rainfall conditions mimicking the natural precipitation in Tuebingen and low rainfall conditions mimicking the precipitation at Madrid (Exposito-Alonso et al., 2019).

The distribution of phenological and fitness traits at each precipitation-location combinations are shown in Figure 2. Significant differences between precipitation-location combinations were observed for fitness and flowering time (*p* < 0.0001). In general, accessions flowered later at Tuebingen compared to Madrid, while low precipitation seemed to delay flowering in both locations indicating that temperature and daylength differences between locations may be the largest driver of differences in flowering time between locations. In general, the accessions exhibited higher fitness in the two high-rainfall treatments compared to low rainfall treatments. Fitness was highest for the high rainfall treatment in Madrid, while the low precipitation treatment at Madrid showed the lowest average fitness. The environment in the high rainfall treatment at Madrid is characterized by simulated rainfall that is similar to the natural precipitation at the common garden location in Tuebingen. Thus, the ample water availability (27.8% SWC) combined with the warm temperatures (median temperature 8.5°C) in Madrid are highly favorable for growth and reproduction in *Arabidopsis*. However, when warm temperatures are combined with inadequate rainfall (16.1% SWC), the overall performance is reduced greatly, as observed for the low average fitness observed in low precipitation in Madrid (M_1_).

**Figure 2.**
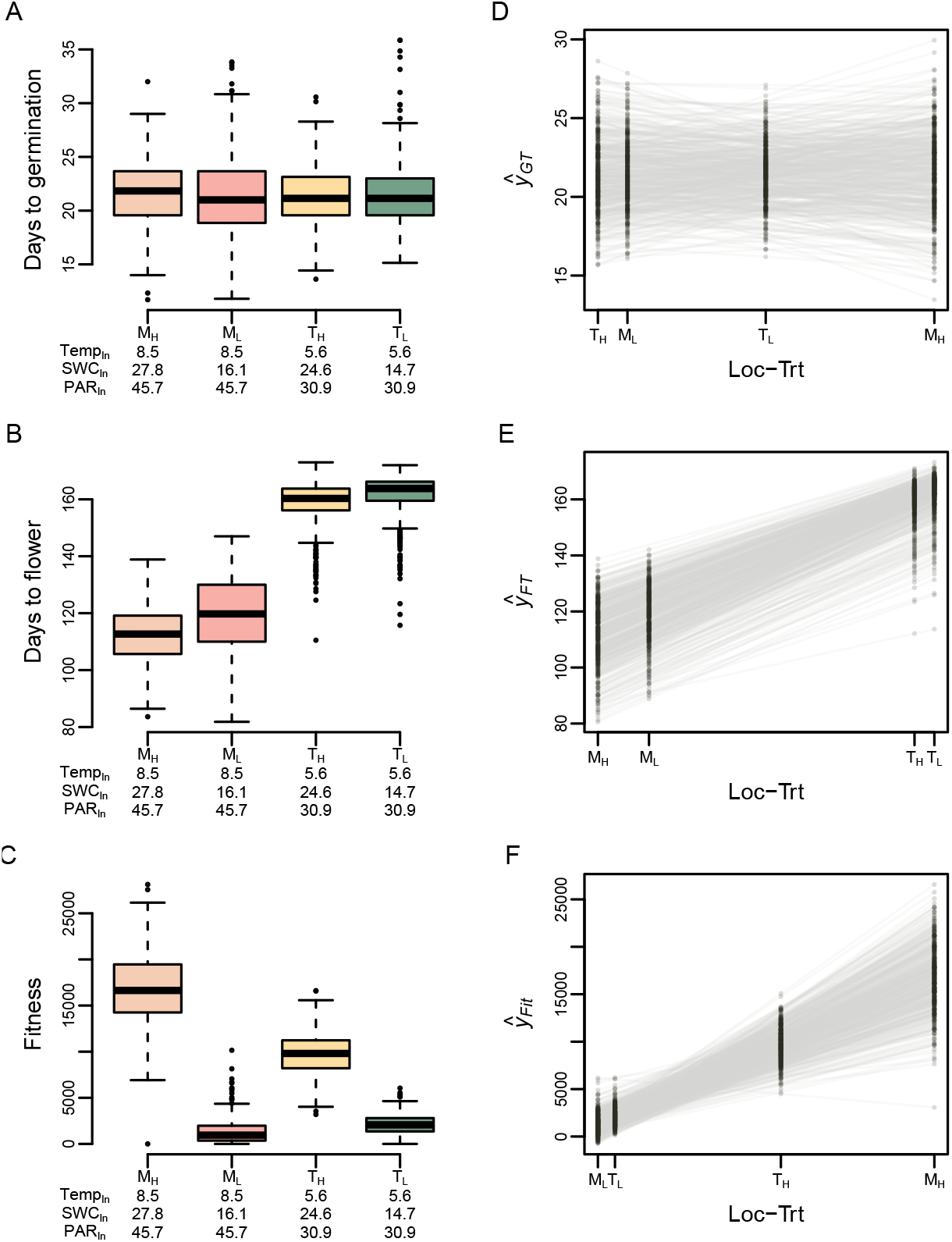
Distribution and plasticity of fitness and phenological traits across contrasting environments. (A-C) Distribution of adjusted means for fitness and phenological traits. Median values for environmental conditions within shelters are shown beneath each boxplot. Temp refers to median temperature in °C, SWC indicates soil water content, and PAR refers to photosythetically active radiation (mol m^−2^ day^−1^). The predicted phenotypic values (ŷ) of each accession in each location-treatment (Loc-Trt) combination is shown in panels D-F and were obtained using the FW approach. ‘M’ refers to common garden in Madrid and ‘T’ indicates common garden in Tuebingen, while the subscripts _*L*_ and _*H*_ refer to the low and high precipitation treatment, respectively.

To estimate environmental plasticity for fitness and phenological traits, we estimated reaction norms for each accession using the FW approach (Finlay and Wilkinson, 1963). Briefly, the FW approach expresses the plasticity for each individual grown across a range of environments as a function of the average population performance at each environment. For each individual, the slope of the linear model expresses the plasticity (or macroenvironmental sensitivity) with respect to average plasticity of the population. The plasticity for each accession with respect to mean performance at each environment is shown in Figure 2D-F.

### Elucidating genetic dependencies between local environmental factors and fitness related traits

To elucidate the genetic interdependencies between local environmental conditions, and fitness and phenological plasticity, we inferred the potential causal genetic relationships between environmental factors and observed phenotypes using four BN structure learning algorithms. Structure learning was performed using the ten latent environmental factors described above and reaction norm slopes for phenological traits and fitness, and the “best” structure was selected based on BIC scores. Of the four algorithms evaluated, the “best” network was given by tabu algorithm (Table 1). Since the primary objective of this study is to elucidate the relationships between local environmental conditions and empirical phenotypes, we focused interpretations of the network on relationships within the Markov blanket for plasticity traits (Figure 3).

**Table 1.** Evaluation of four Bayesian structure learning algorithms. Bayesian network structures were learned using the ten latent environmental variables and plasticity for phenological traits (germination and flowering time) and fitness. The “best” network was selected based on the highest Bayesian information criteria (BIC) and Gaussian BIC values (gBIC). Algo.: algorithm; HC: hill-climbing; MMHC: min-max hill-climbing; RSmax2: general 2-phase restricted maximization

| Algo. | gBIC | BIC |
| --- | --- | --- |
| HC | -1963.02 | -1963.02 |
| tabu | -1962.58 | -1962.58 |
| MMHC | -2480.54 | -2480.54 |
| RSmax2 | -2681.40 | -2681.40 |

**Figure 3.**
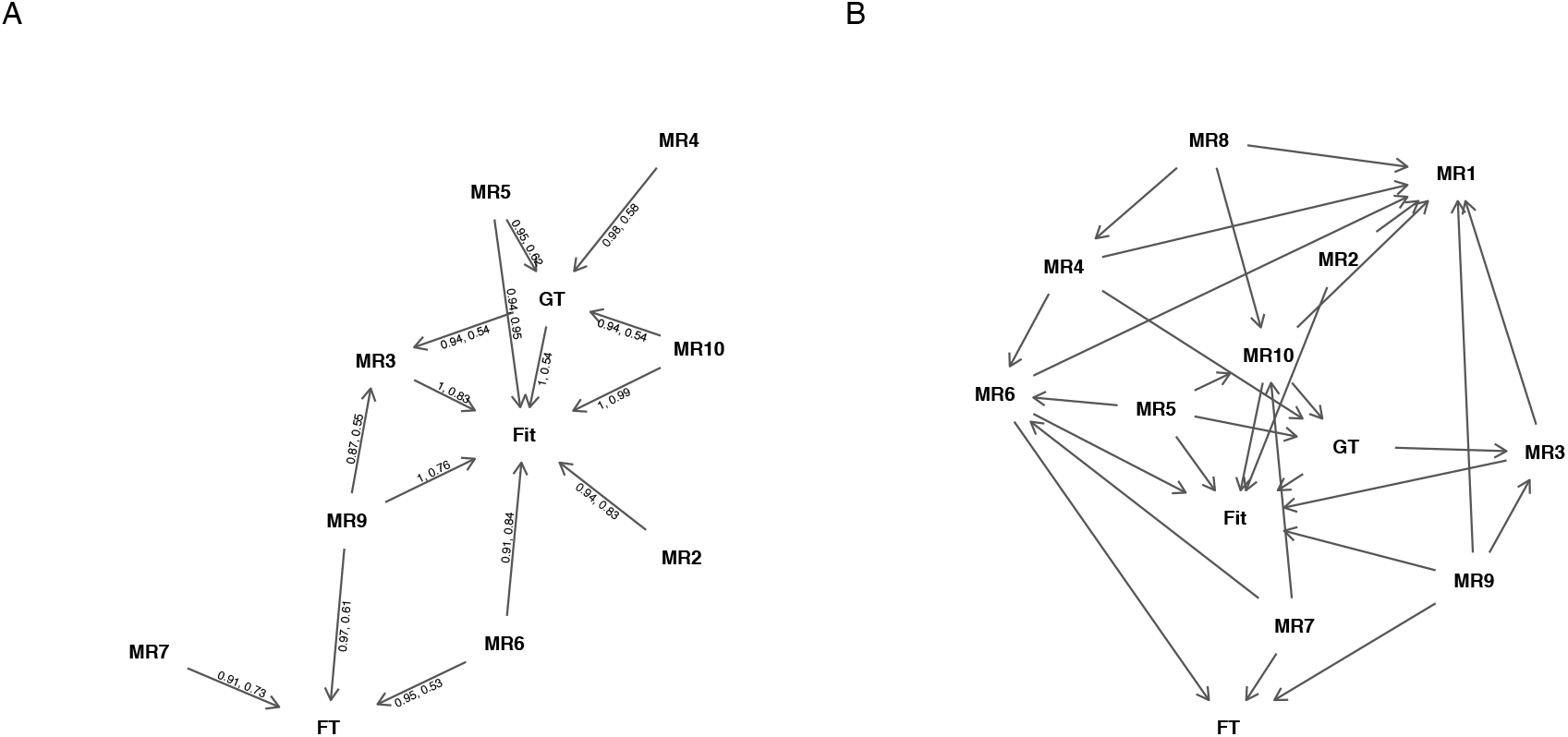
Visual depiction of probabilistic dependancies between environmental variables and empirical phenotypes. The network shown in panel A depicts the Markov blanket for empirical phenotypes and the full network is shown in panel B. A model averaging approach with 500 bootstrap samples was used to learn Bayesian network. The two numbers above each directed edge in panel A shows the proportion of bootstrap samples with the given edge and the proportion of samples with the given direction. The environmental variables are indicated with the “MR” prefix, while the empirical phenotypes are defined as follows: Fit: fitness plasticity; FT: flowering time plasticity; GT: germination time plasticity.

Although the learned structure is complex, several interesting features are apparent. First, of the 29 edges in the network, 41.4% (12 edges) describe relationships from environmental variables to empirical phenotypes, while only 3.45% (1 edge) describe relationships from plastic responses to environmental variables. These results suggest that genomic values for empirical phenotypes are highly dependant on genetic factors associated with adaptation to local environmental conditions. In addition, 51.7% edges (15 edges) were from environmental variables to other environmental variables, and only a single edge was from plastic responses to other plastic responses. Thus, genetic relationships between environmental variables or plastic responses are far more common than relationships from plastic responses to environmental variables.

In addition to overall topological features of the BN, several nodes were identified that were heavily influenced by other variables. For instance, plasticity in flowering time (FT) showed the largest number of indirect effects, suggesting that plasticity in flowering time is highly dependant on genetic effects from adaptation to local environments. A total of seven variables were leading to FT, while three were leading to both plasticity in germination time (GT) and fitness (Fit). Several variables were identified that had indirect effects on many variables. For instance, MR5 and MR9, which describe overall plant productivity, and precipitation and interannual precipitation variability, respectively, each showed indirect effects on four nodes.

### Structural equation modeling

The BN described above represents the probabilistic dependencies between plastic responses and local environmental conditions (Scutari and Denis, 2014). While this approach may provide insights into how variables act on one another, it does not tell *how much* of an effect one variable has on another. To estimate the magnitude of direct (QTL acting directly on focal trait) and indirect (QTL effects transmitted on focal trait by upstream trait) relationships among variables, we performed SEM using the learned structure described above. We leveraged this approach to decompose total genomic values for each environmental variable and empirical phenotype into direct and indirect effects, and examine the covariance between total genomic values and direct genomic values. The matrix of structural equation coefficients is shown in Table 2, and the genomic correlation matrix of direct and total effects is shown in Figure 4.

**Figure 4.**
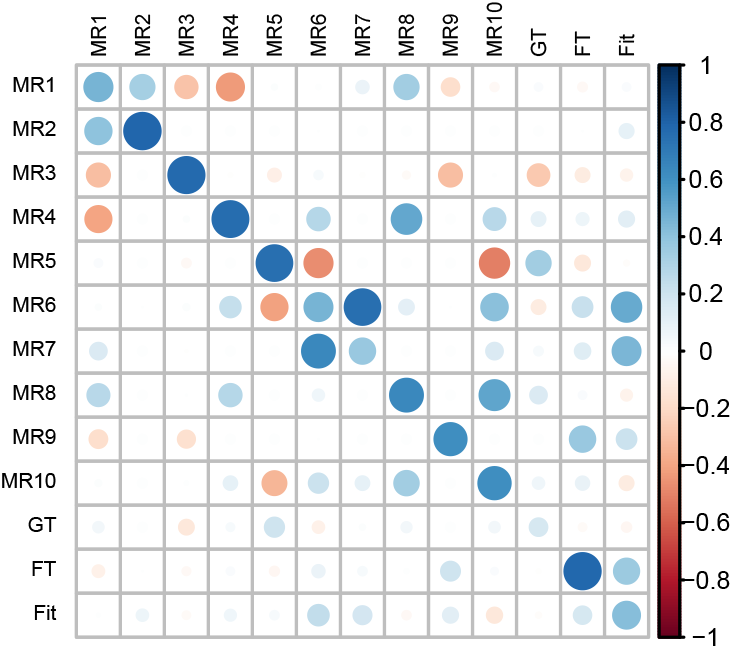
Genomic heritability and correlation for direct and indirect genetic effects. The genomic heritability for total additive genetic effects (*h*^2^) are shown in the diagonal. The upper triangle of the matrix shows the genomic correlation for total effects, while the lower triangle shows the genomic correlation for direct genetic values. Fit: fitness plasticity; FT: plasticity in flowering time; GT: plasticity in germination time

**Table 2.** Structural coefficients estimated using structural equation modeling. Path coefficients for network structure pictured in Figure 3 was estimated using a structural equation modeling approach. The columns indicate upstream nodes, while the rows indicate downstream nodes. Elements with ‘-’ indicate pairs of nodes that are not linked by an edge. Coefficient matrices for structures learned using a Bayesian network approach are typically have zero elements in the diagonal and upper triangle, however the coefficient matrix below has been reordered so that environmental variables are grouped and ordered by name. Fit: fitness plasticity; FT: plasticity in flowering time; GT: plasticity in germination time. Variables with the ‘MR’ prefix indicate latent environmental variables.

|  | MR1 | MR2 | MR3 | MR4 | MR5 | MR6 | MR7 | MR8 | MR9 | MR10 | GT | FT | Fit |
| --- | --- | --- | --- | --- | --- | --- | --- | --- | --- | --- | --- | --- | --- |
| MR1 | - | 0.35 | -0.31 | -0.47 | - | 0.13 | - | 0.6 | -0.23 | -0.18 | - | - | - |
| MR2 | - | - | - | - | - | - | - | - | - | - | - | - | - |
| MR3 | - | - | - | - | - | - | - | - | -0.17 | - | -0.2 | - | - |
| MR4 | - | - | - | - | - | - | - | 0.38 | - | - | - | - | - |
| MR5 | - | - | - | - | - | - | - | - | - | - | - | - | - |
| MR6 | - | - | - | 0.23 | -0.43 | - | 0.81 | - | - | - | - | - | - |
| MR7 | - | - | - | - | - | - | - | - | - | - | - | - | - |
| MR8 | - | - | - | - | - | - | - | - | - | - | - | - | - |
| MR9 | - | - | - | - | - | - | - | - | - | - | - | - | - |
| MR10 | - | - | - | - | -0.33 | - | 0.12 | 0.41 | - | - | - | - | - |
| GT | - | - | - | 0.01 | 0.18 | - | - | - | - | 0.1 | - | - | - |
| FT | - | - | - | - | - | 0.13 | -0.06 | - | 0.31 | - | - | - | - |
| Fit | - | 0.1 | -0.01 | - | 0.13 | 0.34 | - | - | 0.13 | -0.19 | - | 0.11 | - |

The utilization of plastic responses for phenological traits was motivated by several studies that suggest changes in an individual’s life cycle may be an important mechanism for adaptation to specific environmental constraints (Anderson et al., 2012; Vitasse et al., 2013; Augspurger, 2008; Chuine, 2010). While total genomic covariances provide insight into the relationships between total genetic values for two phenotypes, examination of the direct genomic covariances between traits may be more important in the context of the current study, as the covariance of direct genomic effects is driven by QTL that have an effect on both environmental adaptation and plasticity or QTL that affect each trait independently but are in tight LD (Valente et al., 2013). For direct genomic effects, the strongest positive genomic correlation between plastic responses and environmental variables was observed for Fit and MR6 (*r_gdirect_* = 0.24), which is a composite of temperature conditioning indices with lower values indicate a potential for high temperature stress on vegetative biomass. Fit also showed positive direct genomic correlation with MR7 (*r_gdirect_* = 0.18), a variable composed of indices quantifying plant health, and MR9 (*r_gdirect_* = 0.13), which quantifies precipitation and interannual variability in precipitation. Collectively, these results indicate that the accessions that harbor alleles for reduced sensitivity of fitness to temperature gradients likely also harbor alleles associated with adaptation to warm, low rainfall environments.

In addition to Fit, relatively strong positive direct genomic correlation was observed between FT and MR9 (*r_gdirect_* = 0.20), as well as GT and MR5 (*r_gdirect_* = 0.21). However, the slope for FT largely represents the sensitivity of flowering time to differences in photoperiod and/or temperature for an accession, with lower values indicating more similar flowering times between common garden locations.

Therefore, it is unclear whether the non-zero direct genomic covariance between these variables indicates a common mechanism, or potential confounding of photoperiod insensitive accessions originating from more southern locations.

## Discussion

Environment association analyses have become popular approaches to elucidate the genetic basis of local adaptation in the absence of fitness measurements in multilocation common garden trials (Fournier-Level et al., 2011; Yoder et al., 2014; Lasky et al., 2015). The aim of EAA is to identify genes or loci that may impact traits that confer fitness along an environmental gradient. However, when fitness is measured in multiple common garden locations along an environmental gradient, the change in fitness as a function of mean population performance provides a single metric that describes the impact of the environment on fitness. Moreover, when this metric is introduced as the response variable in genome-wide association studies, strong associations indicate the presence of gene(s) that may influence fitness along the environmental gradient.

In the current study, we seek to integrate both data types in the SEM framework to examine the genetic interdependencies and covariances between changes in fitness and phenology in multiple environments and local environmental conditions. However, whereas most EAA estimate the effects of individual loci, we predict the total genetic values (i.e. the summation of QTL effects for a given genotype) for each variable. Thus, in cases where collection sites and common garden locations follow the same gradients, we expect covariance in genetic signals that impact both variables directly. Consistent with this expectation, we observed non-zero genetic covariance between local environmental conditions and changes in fitness between common garden locations. For instance, Fit showed positive correlation of direct genetic effects for MR6, as well as MR7. The latent variables MR6 and MR7, capture the favorably of the local environment for plant growth. Thus, higher values indicate environments that have favorable conditions for plant growth and, on a whole, are highly productive. Moreover, Fit describes the changes in fitness driven largely by water availability, with higher values indicating greater fitness in high-rainfall treatment in Madrid and low values indicating low fitness in low-rainfall treatment in Madrid (Figure 2). Thus, the positive genomic correlation of direct effects indicates that accessions harboring alleles for high fitness in simulated, high-productivity environments will also tend to harbor alleles associated with adaptation to highly productive local environments. Although weaker than the direct genomic correlation for MR6 and MR7, Fit also showed positive genomic correlation with a latent environmental variable that largely captured precipitation and precipitation variability of the local environment, with higher values indicating higher precipitation (MR9; *r_gdirect_* = 0.13). Collectively, these results indicate that fitness in response to some local environmental conditions may be regulated common genetic mechanisms that affect fitness in simulated environments. However in either case (e.g., local environment associations or common garden fitness), the traits that impact fitness are largely unknown.

Phenotypic plasticity is an important process that allow plants to quickly modify physiology, morphology, or phenology in response to changes in the environment (Bradshaw, 1965). Individuals that exhibit greater plasticity may be better positioned to respond to new environmental constraints, as novel phenotypes bought on by environmental change may provide persistence in the short-term (West-Eberhard, 2005; Matesanz et al., 2010; Nicotra et al., 2010; Valladares et al., 2014). However, phenotypic plasticity is not always advantageous (DeWitt et al., 1998; Ghalambor et al., 2007). For instance, Scheepens and Stöcklin (2013) showed that increased temperature leads to early flowering, but reduced seed set in *Campanula thyrsoides*. Thus, it is important to couple observations of plasticity across an environmental gradient with measurements of fitness in the same environments to determine whether phenotypic plasticity can be a mechanism underlying fitness. Here, we utilized measures of fitness and empirical phenotypes in four environments. Correlations for reaction norms for fitness and phenological traits showed a significant, albeit weak, correlations between Fit and GT (*r* = −0.09, *p* = 0.417) and Fit and FT (*r* = 0.18, *p* < 0.0001), indicating that these changes in fitness are associated with changes in phenology. In the case of FT, the positive correlation indicates that accessions that show greater plasticity in flowering time (positive slope for FT meaning delayed flowering in Germany relative to Madrid) tend to exhibit greater fitness in high-precipitation regimes relative to low-precipitation regimes (i.e., positive slope for Fit). Correlation provides a simple means to measure the relationships between two traits within a population. However, a non-zero correlation does not necessarily indicate that the outcome/expression for one characteristic is dependant on another. BN approaches on the other hand, have been developed to elucidate probabilistic dependencies among a group of interrelated variables (Pearl, 2014; Scutari and Denis, 2014). The BN shown in Figure 3 shows an directed edge from GT to Fit, indicating that changes in fitness across the common garden environments is dependant on changes in germination time. However, no edges were found between FT and Fit, indicating that although these two characteristics covary, changes in Fit may not be dependant on changes in FT.

Although it is seemingly a natural tendency to view these dependencies as causal relationships, it is important not to over interpret results from BN. While BN are a powerful approach to assess the interdependencies between variables, structure learning with BN imposes several constraints that may limit its applications in biology. One major limitation is that BN do not allow feedback loops or cyclical relationships in the structure, which are pervasive throughout biology especially at a molecular level (Scutari and Denis, 2014). Thus, if the underlying causal relationships between traits involves feedback loops, the structure learned with BN will likely be inaccurate (Valente et al., 2013). Thus, the network might reflect highly probable relationships between variables, but may not represent the true causal relationships that give rise to the data. Secondly, in the current study, BN were constructed using a mixture of observational and experimental data. In the absence of randomization, dependencies observed in Bayesian networks constructed using observational data may be driven by unobserved confounders, thereby making causal claims based on the data problematic (Bello et al., 2018, see for review). Nevertheless, causal relationships can be learned from the data and should be used to generate hypothesis for further studies. In our study, BN revealed dependencies between plasticity in fitness and several environmental variables. Fitness in a given environment is largely the consequence of a trait or traits that confer adaptation to a set of environmental conditions. In other words, fitness is not a mechanism for local adaptation, but rather is a measure of adaptation. Thus, we expect that fitness in a given location/precipitation regime should be highly dependant on mechanisms that were selected by environmental pressures in the accessions’ local environments, and this expectation is largely confirmed by the network learned from the data (Figure 3). However, covariance in direct effects for other variables, such as between FT and MR9, may not be so easy to explain. The latent environmental variable MR9 largely captures precipitation and precipitation variability, as the manifest variables spring precipitation, precipitation of the wettest month, and interannual precipitation variability load onto MR9. The positive direct covariance between MR9 and FT suggest that accessions that harbor alleles associated with adaptation to environments with high precipitation will also tend to harbor alleles associated with higher plasticity in flowering time. However, plasticity in flowering time is largely driven by differences in day length and temperature between common garden locations rather than by precipitation regimes (Figure 2). Thus, it is questionable whether the direct genomic covariance is due to QTL that affect adaptation to precipitation gradients and the sensitivity of flowering time to photoperiod and/or temperature, or if this is driven by unaccounted, confounding effects within the data. Projection of phenotypic values for FT on collection sites show clusters of accessions originating from the Northern Iberian peninsula and Southern Sweden with low plasticity for flowering time (Supplemental Figure S4). Moreover, these regions also exhibit low values for MR9. Further studies or alternative experimental designs are necessary to determine whether this covariance is due to common effects on adaptation to precipitation gradients and plasticity in FT, or are due to sampling bias. Thus, while BN can provide important insight into the interrelationships between traits, when these networks are constructed using observational data we should view these results with caution rather than to discount inferred relationships as spurious.

While BN describe the probabilistic dependencies among variables, they only provide insight into the structure of relationships in the data. In many cases, we are interested in understanding how genetic effects for an upstream trait affect the outcome of a downstream trait. SEM provides a means to estimate path coefficients according to a predefined network structure, as well as partition phenotypic values into genetic values that affect a trait directly (i.e., direct genetic values) and genetic values that are due to genetic effects acting directly on upstream variables (Gianola and Sorensen, 2004; Valente et al., 2013). In some sense, estimates of the structural coefficients may seem like the most attractive component of SEM, as these describe how intervention on an upstream variable (e.g., a latent environmental variable) will impact the outcome of the downstream variable (e.g., empirical phenotype) given the direct effects for the downstream variable remain unchanged (Gianola and Sorensen, 2004). However in the current study, we have data that is a combination of latent environmental variables and empirical phenotypes. Thus, a more biologically meaningful question is whether QTL that have a direct effect on adaptation to an environmental gradient also have a direct impact on some observable phenotype. Non-zero covariance in direct effects between local environmental conditions indicates the presence of common QTL, or independent QTL that are tightly linked (Valente et al., 2013). Thus, identification of such QTL can provide important insights into the common mechanisms that impact adaptation to local environments and plasticity.

## Acknowledgements

Funding for this research was provided by the National Science Foundation (United States) through Award No. 1736192 to GM.

## Supplemental Materials

- **Supplemental Figure S1.** Geographic locations for all 1,035 *Arabidopsis* accessions.
- **Supplemental Figure S2.** Heatmap for 55 manifest environmental variables.
- **Supplemental Figure S3.** Projection of phenotypic values for flowering time plasticity on collection sites for 515 accessions.
- **Supplemental File S1.** Factor loading from exploratory factor analysis.
- **Supplemental File S2.** Description of latent and manifest variables, and their loadings from confirmatory factor analysis.

## Supplemental Data

**Figure S1.**
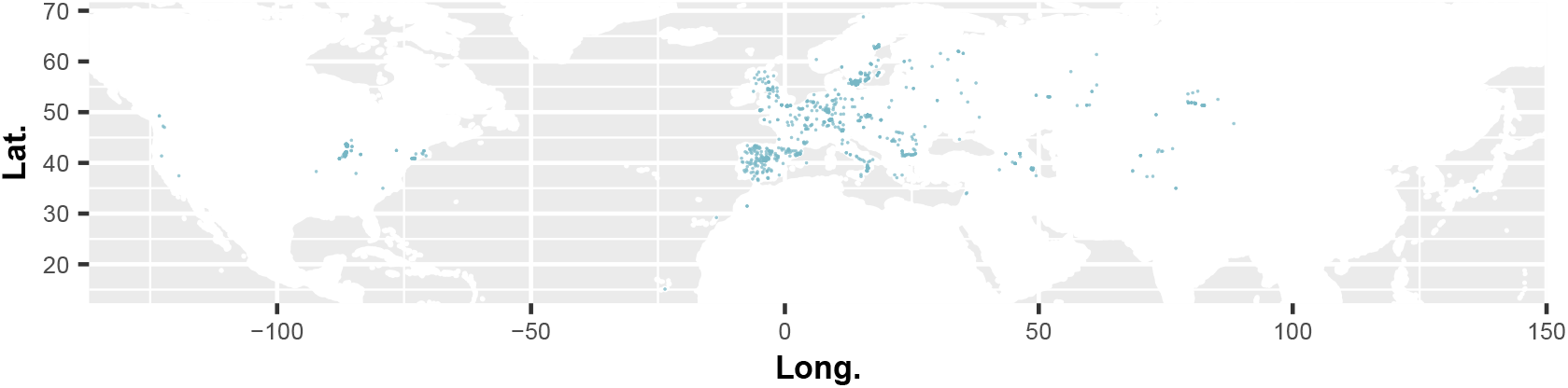
Geographic locations for all 1,035 *Arabidopsis* accessions. The locations for 1,035 accessions used to define latent environmental variables.

**Figure S2.**
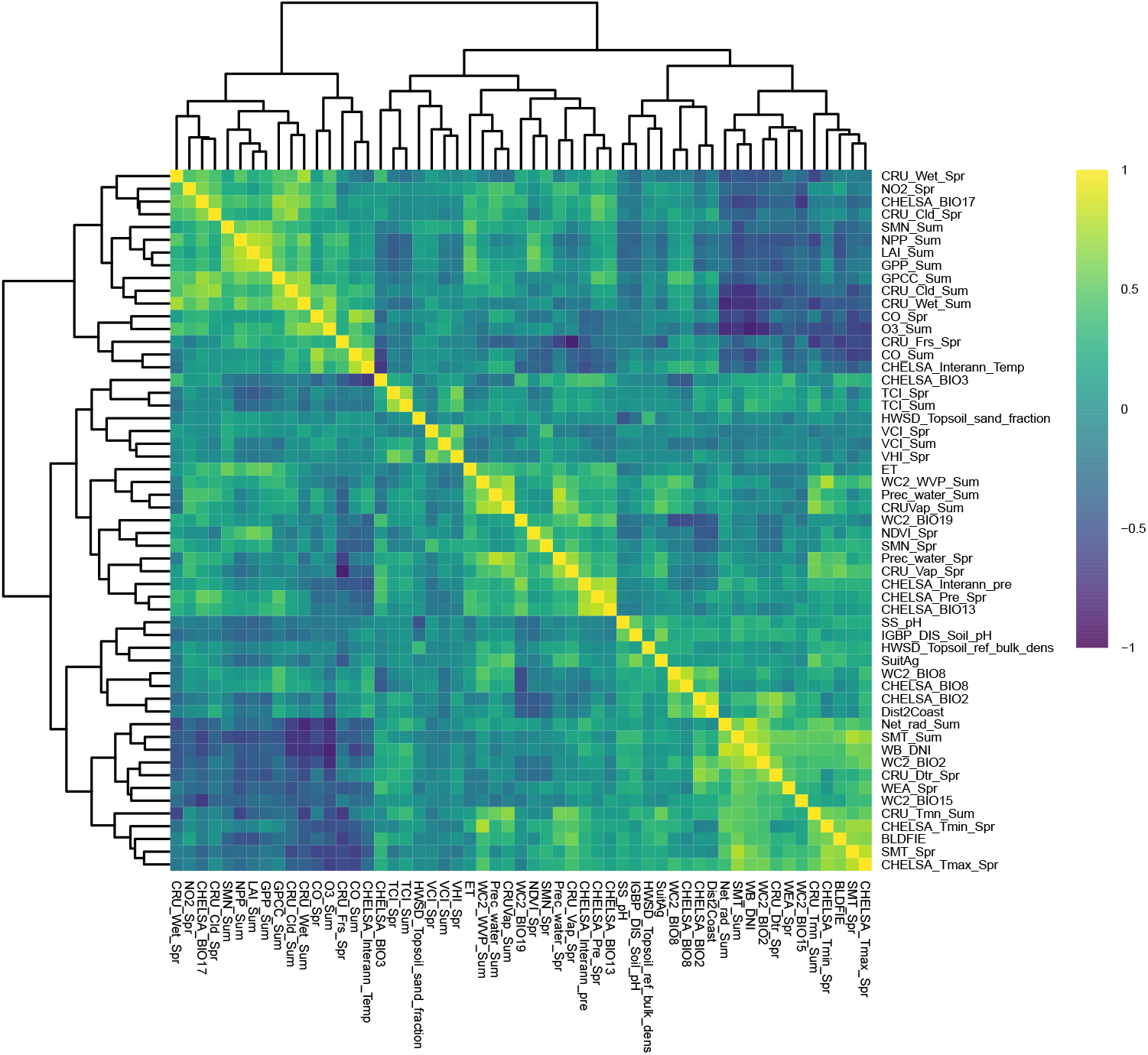
Heatmap for 55 manifest environmental variables. Spearman’s method was used to generate the correlation matrix.

**Figure S3.**
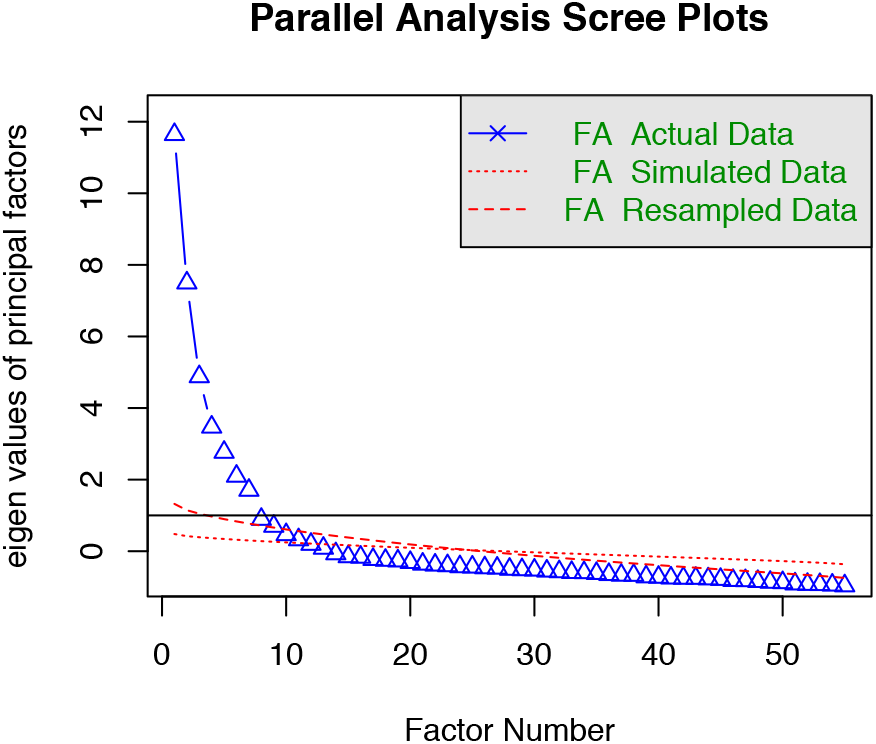
Scree plot indicting the optimal number of latent factors for 55 environmental variables. Parallel factor analysis was performed using the `psych` package in R. This approach generates scree plots for the observed data and compares the results with scree plots generated from a random data matrix of the same size as the observed data set.

**Figure S4.**
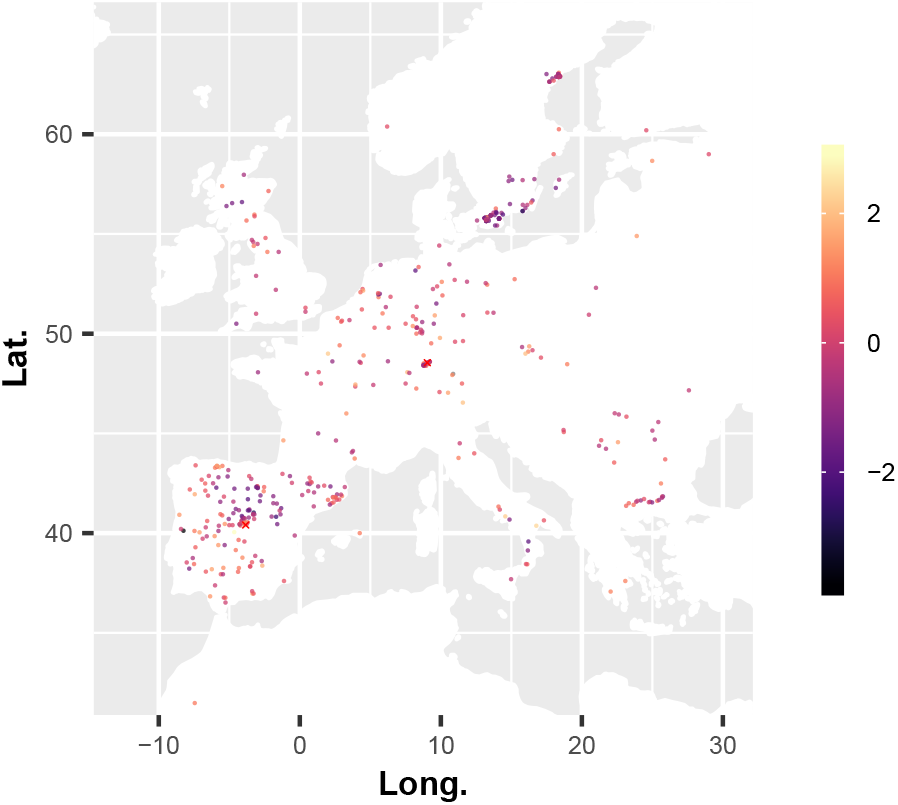
Projection of phenotypic values for flowering time plasticity on collection sites for 515 accessions. Higher plasticity values indicate a greater delay in flowering time Tuebingen relative to Madrid, and is indicated by the continuous color scale on the right. The red ‘X’ indicates the two common garden locations.

